# Regional dynamics of fractal dimension of the left ventricular endocardium from cine computed tomography images

**DOI:** 10.1101/791731

**Authors:** Ashish Manohar, Lorenzo Rossini, Gabrielle Colvert, Davis M. Vigneault, Francisco Contijoch, Marcus Y. Chen, Juan C. del Alamo, Elliot R. McVeigh

## Abstract

We present a method to leverage the high fidelity of CT to quantify regional left ventricular function using topography variation of the endocardium as a surrogate measure of strain. 4DCT images of 10 normal and 10 abnormal subjects, acquired with standard clinical protocols, were used. The topography of the endocardium was characterized by its regional values of fractal dimension (F_D_), computed using a box-counting algorithm developed in-house. The average F_D_ in each of the 16 American Heart Association segments was calculated for each subject as a function of time over the cardiac cycle. The normal subjects showed a peak systolic percentage change in F_D_ of 5.9% ± 2% in all free-wall segments, while the abnormal cohort experienced a change of 2% ± 1.2% (p < 0.00001). Septal segments, being smooth, did not undergo large changes in F_D_. Additionally, a principal component analysis was performed on the temporal profiles of F_D_ to highlight the possibility for unsupervised classification of normal and abnormal function. The method developed is free from manual contouring and does not require any feature tracking or registration algorithms. The F_D_ values in the free wall segments correlated well with radial strain and with endocardial regional shortening measurements.

## 1. Introduction

Over the past three decades, significant effort has been focused on developing robust and accurate methods for quantitative evaluation of regional cardiac function which has implications in the diagnosis, treatment, and follow-up of patients with cardiac diseases such as myocardial ischemia^1,2^, heart failure^3^, cardiotoxicity^4^, and dyssynchrony^5,6^. Left ventricular (LV) ejection fraction (EF), global longitudinal strain (GLS)^7^, and other global metrics have been useful in the diagnosis of heart disease; however, significant drops in EF are typically detectable only in advanced disease stages, and GLS is not a local measurement. Additionally, regional dysfunction often precedes global dysfunction and global metrics fail to identify specific regions of abnormally functioning cardiac tissue.

Recent advances in x-ray computed tomography (CT) technology have led to very short scan times (∼140 ms for a full high resolution 3D volume of the heart), permitting measurement of 3D motion of the whole heart for a full cardiac cycle within a single heartbeat acquisition^8–12^. Additionally, the superior spatial resolution of CT has made it possible to visualize and track fine endocardial features such as trabeculae carneae^13^. Viewing a simple surface rendered LV blood volume as a 3D movie shows directly that the trabeculae undergo significant deformation during systolic contraction. The aim of this work was to exploit the high fidelity of x-ray CT to derive a simple metric of regional cardiac function with minimal operator involvement and fast post-processing of the images. We hypothesized that the variation of LV endocardial topography across the cardiac cycle, characterized by regional values of its surface fractal dimension (F_D_), could be obtained from simple threshold data and can serve as a surrogate measure of regional strain in those regions that have trabeculation. Other shape-based measures for cardiac motion analysis have been proposed and established. McEachen & Duncan, 1997^14^ established a technique to quantify the non-rigid and non-uniform motion of the LV by matching local segments of the endocardial contour across time using a shape-based optimization function. Duncan et al., 1988^15^ proposed a shape-based model using the bending energy required to deform LV endocardial contours from an undeformed ‘normal’ state to their individual characteristic shape, as a method to quantify abnormal variations in the end-diastolic shape of the LV, which are suggestive of infarcts and LV remodeling.

The fractal dimension of a surface like the endocardium is a quantitative measure of its geometric complexity^16^. A smooth surface has a dimension of 2 but surfaces with intricate topography can have higher fractal dimensions in the range of *2<F*_*D*_*<3*, as they become increasingly volume-filling. Figure 1 illustrates the relationship between the topographical complexity of a surface and its fractal dimension. This figure was created using the Diamond-Square algorithm for generating fractal surfaces via a fractional Brownian process^17^. Different surfaces were obtained for different values of the Hurst exponent *H*, which prescribes the fractal dimension as *F*_*D*_ *= 3 – H*^16^. Figure 1A shows a gently warped square patch with *F*_*D*_ *= 2.1*. Figures 1B and 1C have increasing amounts of roughness resulting in fractal dimensions of 2.5 and 2.8 respectively.

**Fig. 1.**
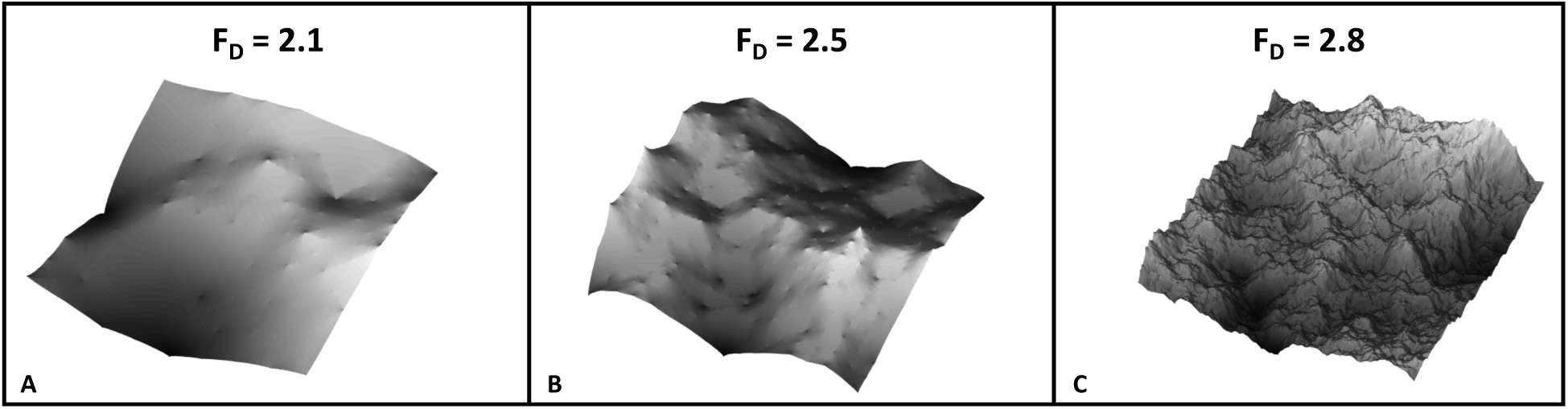
Illustration of the relationship between fractal dimension (F_D_) and topography. (**A**-**C**) Varying amounts of topography corresponding to Hurst exponents of 0.9, 0.5, and 0.2 added to a regular square, resulting in F_D_ of 2.1, 2.5, and 2.8 respectively.

Previously, Moore & Dasi, 2013^18^ measured the global fractal dimension of the LV endocardium across the cardiac cycle from cine CT images. They reported that the endocardial trabeculae contract during systolic ejection, giving the endocardium its lowest dimension at end-systole (global *F*_*D*_ *= 2.02*) followed by their relaxation and subsequently the highest dimension of the endocardium at end-diastole (global *F*_*D*_ *= 2.2*). The difference between the end-diastolic and the end-systolic global F_D_ values were found to be statistically significant (p < 0.003), suggesting that changes in endocardial F_D_ across the cardiac cycle are in accordance with the motion of the LV wall. While this study provided useful insight in hypothesizing the role of the trabeculae in ejection, there was no measurement of regional values of endocardial F_D_ nor a correlation of F_D_ to regional myocardial function. Additionally, Captur et al., 2013^19^ used fractal analysis on short-axis CMR images of 135 individuals and reported that patients suffering from LV non-compaction (LVNC) had significantly higher values of endocardial F_D_ (p *<* 0.00001). While the study provided a new quantitative and reproducible estimate of LVNC, there was again no regional analysis nor did they relate the F_D_ to regional myocardial function. Additionally, they computed a global F_D_ for 2D short axis MR slices, which require multiple heartbeats and are subject to through plane motion. In this work, we quantify regional left ventricular function in both normal and abnormal left ventricles from true 3D high-resolution cine CT images acquired within a single heartbeat, and compare the regional F_D_ estimates to radial strain and endocardial regional shortening measurements.

## 2. Methods

### 2.1 Subjects

For this study, 4DCT scans of 20 human subjects (4 female, 16 male) acquired from two centers (The National Institutes of Health (NIH), Bethesda, Maryland and the University of California San Diego (UC San Diego), California) were used. All subjects were scanned under IRB approved protocols at the two centers and each scan sampled the entire cardiac cycle of one heartbeat. Subjects were categorized to have normal function if the following three conditions were satisfied:

1. Ejection fraction >60% as measured from CT.
2. No wall motion abnormalities observed on the cine CT images.
3. Peak end-systolic radial strain >50%^20^ as measured from the CT images using the Aquarius iNtuition (TeraRecon Inc.) analysis package, shown in Fig. 2.

Based on the above, 10 of the 20 subjects were categorized to have normal LV function; the remaining 10 were categorized as abnormal. Additionally, all abnormal subjects had an expert radiologist and cardiologist clinical assessment. The normal and abnormal subjects had ages of (mean ± SD) 63 ± 15 years and 63 ± 19 years respectively.

**Fig. 2.**
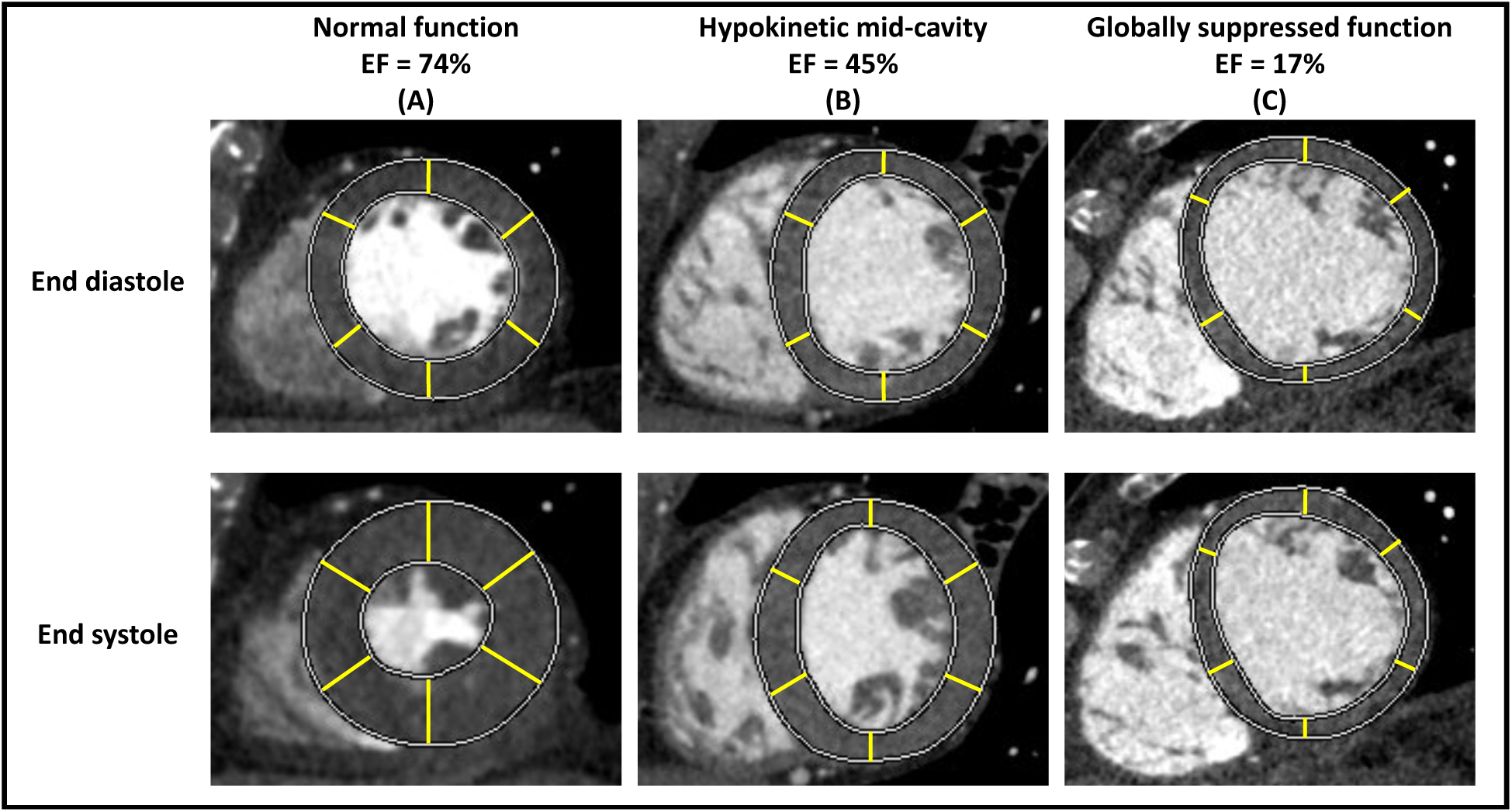
Radial strain measurements by traditional contouring methods in three sample subjects. The top panel shows mid-cavity short axis slices at end-diastole and the bottom panel shows mid-cavity short axis slices at end-systole; both panels are in the traditional AHA orientation. The epicardial and the endocardial contours were automatically generated using the Aquarius iNtuition (TeraRecon Inc.) analysis package with necessary user intervention to manually correct identified contours. The yellow rays are visual aids representing the wall thickening in each of the 6 traditional AHA segments. (**A**) Subject with normal function and an EF of 73%. (**B**) Subject with a hypokinetic mid-cavity and an EF of 45%. (**C**) Subject with global dysfunction and an EF of 17%. AHA: American Heart Association, EF: ejection fraction.

### 2.2 CT Imaging

The CT images were acquired using standard imaging protocols prescribed by the two clinical centers. The 20 subjects were scanned using two different scanner models: a 320 detector-row (Aquilion ONE, Canon Medical, Otawara, Japan) at NIH and UC San Diego and a 256 detector-row (Revolution CT, General Electric Medical Systems, Wisconsin) at UC San Diego, both of which enabled complete coverage of the heart from a single table position. The images were acquired using an electrocardiogram (ECG) gated protocol with inspiratory breath-hold. Optimal opacification from contrast dye was achieved using real-time bolus tracking. For each image, 70-100 ml (depending on patient weight) of Iopamidol (Isovue 370, Bracco) was injected at 5 ml per second with a 40 ml saline flush.

Dose modulation across the cardiac cycle was not used in any of the 20 subjects in order to avoid a temporal change in image quality due only to changing tube current. In this way, the temporal evolution of fractal dimension was used as a test of the reproducibility of the calculation. A region of interest (ROI) was drawn inside the LV blood pool to estimate the signal-to-noise ratio (SNR), which was calculated as the ratio of the mean to the standard deviation of the Hounsfield Units (HU) within the ROI. The mean SNR values found in the normal and abnormal subjects were (mean ± SD) 14.5 ± 1.5 and 14 ± 3 respectively.

The images were reconstructed using each manufacturer’s standard reconstruction algorithm as implemented clinically. The scans were reconstructed into images of 512 × 512 pixels with in-plane resolutions between 0.32mm and 0.48mm and between 100 to 320 slices with resolutions of 0.5mm to 1mm in the *z*-direction. Reconstruction was performed at regular intervals of the R-R phase of the ECG, with 5% and 10% being the minimum and maximum intervals. Neither subject enrolment nor image reconstruction was performed for the purpose of our particular study. The images used in this work were directly obtained from the clinic; hence, were reconstructed with different fields of view and slice thicknesses as determined by the clinical protocol used for that study.

### 2.3 Segmentation of the LV Endocardium

Image processing was performed using in-house command scripts developed in MATLAB (Mathworks Inc.), unless specified otherwise. All images were input into the analysis software as anonymized DICOM files. Only standard, well characterized image processing steps were used in the segmentation process in order to highlight the fractal dimension measurements; not a novel segmentation technique. A summary of the process is shown in Fig. 3. The main steps were:

**Fig. 3.**
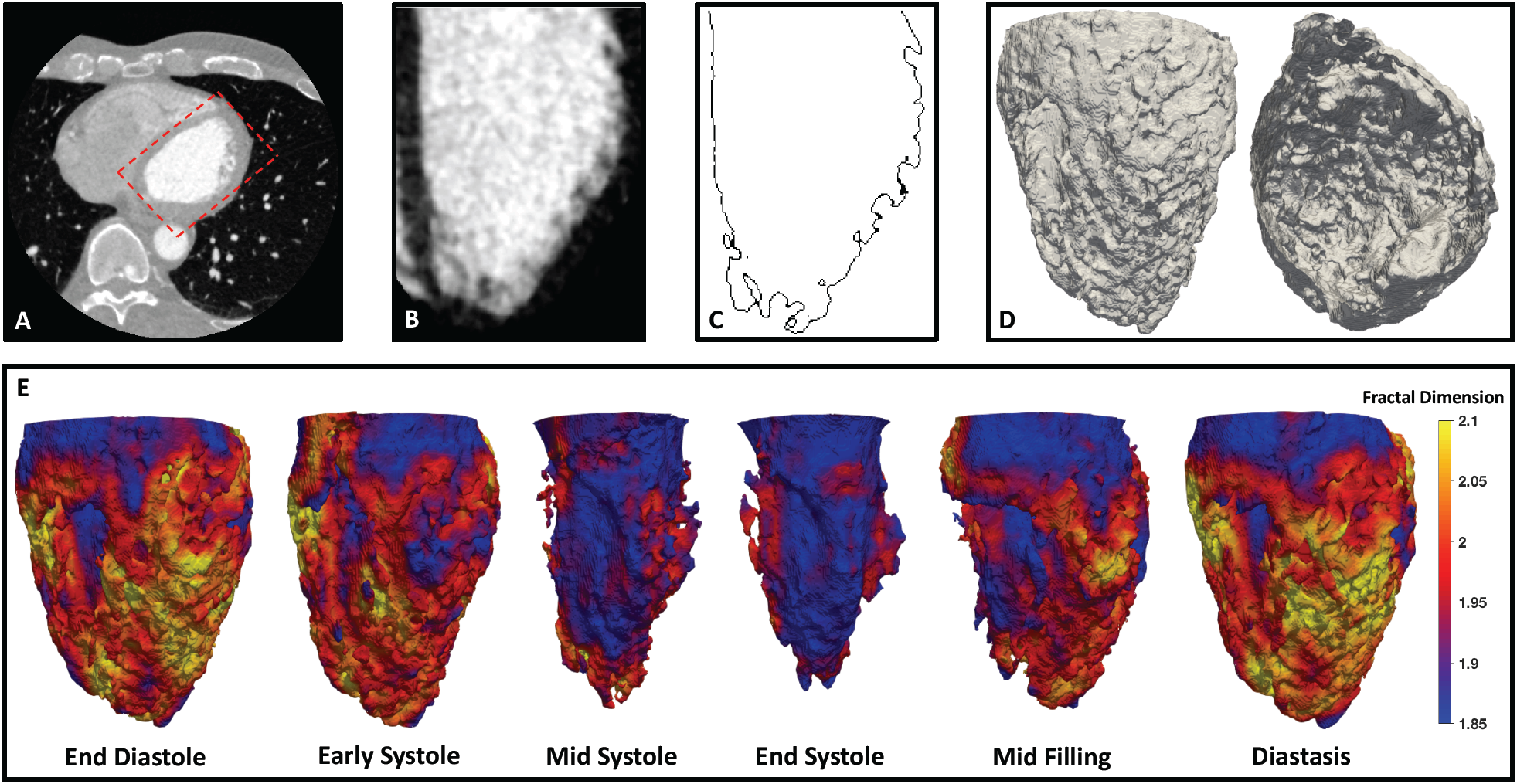
Steps of the proposed method. (**A**) Axial slice of a CT image with the LV defined by the dashed red box. (**B**) Long axis slice of the LV blood pool looking at the anterior wall with the inferior wall hidden from view, the septum on the left, and the lateral wall on the right. (**C**) Long axis slice of the LV endocardium (same view angle as **3B**) extracted from the binarized LV blood pool. (**D**) 3D surface renderings of the LV endocardium showing topography of trabeculae carneae. The left view is of the free wall of the endocardium, with the anterior wall on the left, the inferior wall on the right, and the septum hidden from view. The right view shows the interior of the LV cavity to highlight the texture of the trabeculae, looking from the base down into the apex with the inferior wall at 12:00, the lateral wall at 3:00, the anterior wall at 6:00, and the septum at 9:00. (**E**) 3D surface renderings of the LV endocardium with its regional F_D_ maps across 6 cardiac phases, looking at the free wall of the LV with the septum hidden from view, the anterior wall on the left and the inferior wall on the right. During systolic contraction, the trabeculae collapse inwards upon each other, giving rise to a smoother topography of the endocardium and hence to lower values of F_D_.

1. All images were resampled to an isotropic resolution of 0.3 × 0.3 × 0.3 mm^3^. This resolution was chosen based on the highest native resolution of our images to avoid aliasing effects by down-sampling to a lower resolution.
2. The LV blood pool was segmented and binarized by applying a suitable threshold between 250 and 700 HU, which was set by the user after estimating the mean and standard deviation of the HU within the blood pool according to the following equation 

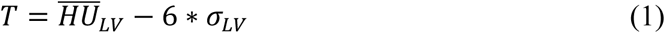

where *T* is the input threshold, 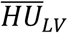 is mean signal of the blood in a 4000 pixel ROI within the blood pool, and *σ*_*LV*_ is the standard deviation around that mean.
3. The coronaries and any remnants of the right ventricle (RV) were automatically pruned by retaining the single largest connected region.
4. The endocardium was extracted from the binary LV blood pool using a 3D discrete Laplacian filter^21^.

### 2.4 Calculation of Fractal Dimension

The calculation of F_D_ was performed using in-house code developed in the FORTRAN programming language. Regional characterization was achieved by calculating the F_D_ of the LV endocardium contained within local neighborhoods (subsets) of 32 × 32 × 32 voxels, each of which corresponded to a volume of 9.6 × 9.6 × 9.6 mm^3^. A box-counting algorithm was used to compute F_D_ as^16^

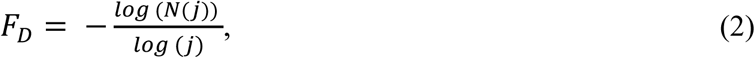

where *N(j)* is the number of boxes containing non-zero voxels within a grid defined by cubic boxes of size (i.e. edge length) *j*. The entire LV endocardium was characterized by moving the local neighborhood by 8 voxels sequentially in the *x, y*, and *z* directions. For each neighborhood, the box size varied between *j = 1* and *j = 16* and F_D_ was obtained by linear regression of *log(N(j))* with respect to *log(j)*. The computed regional F_D_ values were linearly interpolated back onto every voxel of the LV endocardium, completing its regional characterization as shown in Fig. 3E. The box-counting algorithm was validated using synthetic standard fractals of known F_D_, obtaining excellent agreement (Appendix A).

### 2.5 Calculation of Average Fractal Dimension in 16 AHA Segments

The endocardium at each time frame was divided into the 16 standard American Heart Association (AHA) segments^22^ and the F_D_ of each segment was calculated as the arithmetic mean of all voxel values within that segment. The temporal evolution of *F*_*D*_*(t,s)* for each segment, *s*, was mapped onto 20 time points, *t*, across the R-R interval from 0% through 95% and was smoothed using a low-pass filter (first three Fourier harmonics).

### 2.6 Correlation between Peak Systolic Radial Strain and Peak Systolic Change in Regional F_D_

Peak end-systolic radial strain (ε_rr_) measurements were acquired in all 16 AHA segments for all 20 subjects used in the study. The measurements were made using the Aquarius iNtuition (TeraRecon Inc.) analysis package by drawing epicardial and endocardial contours at end-diastole and end-systole as shown in Fig. 2. From these contours, percent radial thickening (radial strain) was calculated for each of the 16 AHA segments. Although the analysis package generated automatic epicardial and endocardial contours, user intervention was often required to manually correct identified contours. Peak systolic change in F_D_ (ΔF_D_) was also calculated in all 16 AHA segments for all subjects and ε_rr_ was plotted against ΔF_D_ as a scatter plot.

### 2.7 Correlation between Temporal Changes in F_D_ and Temporal Changes in Endocardial Regional Shortening (RS)

In order to evaluate if the temporal changes in regional F_D_ correlated with changes in LV endocardial regional shortening (RS) across the cardiac cycle, we compared results from the F_D_ analysis with results from the recently developed SQUEEZ algorithm^13^ on the same set of images. SQUEEZ has been shown to correlate well with MRI tagging derived circumferential shortening^23^. Our use of SQUEEZ to derive RS in this study was driven by the fact that the two metrics (F_D_ and SQUEEZ) were derived from the same images. This eliminated the problem of attempting to measure strain in hearts at different times (as is the case for comparisons between different imaging techniques such as CT vs echocardiography or CT vs MRI). The correlation with RS was performed as a comparison, not as a validation because RS derived from CT has not yet been accepted as a “gold standard” for endocardial strain.

SQUEEZ is a measure of endocardial 2D strain (it is the geometric mean of circumferential and longitudinal strain – it does not measure radial strain). Regional Shortening (RS = SQUEEZ-1) is measured by tracking features on the endocardium from 4DCT data and is computed as the square root of the ratio of areas of corresponding patches between a template mesh and a target mesh according to the equation 

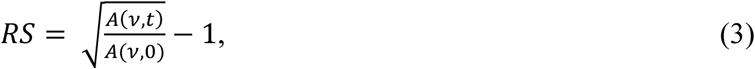

where *A*(*v, t*) is the area of a patch *v*at time *t* and *A*(*v*, 0) is the area of the same patch at time *t* = 0 (end-diastole).

A principal component analysis (PCA) was performed on the temporal profiles of F_D_ and RS across the cardiac cycle for all AHA segments. This was done for two reasons: 1) to evaluate the possibility of achieving an unsupervised classification of normal and abnormal function, and as a comparison between F_D_ and RS. The PCA was performed independently for both metrics by pooling all subjects together with no prior labeling. For simplicity, we kept the first two PCA modes for both metrics and approximated *F*_*D*_(*t*) and *RS*(*t*) as 

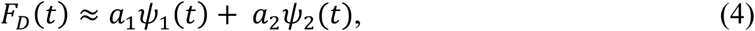

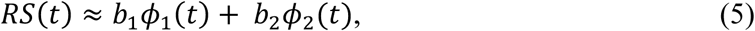

where the first modes *ψ*_1_(*t*)/ *ϕ*_1_(*t*) and the second modes *ψ*_2_(*t*)/ *ϕ*_2_(*t*) capture patterns of slow and fast wall motion respectively.

## 3. Results

### 3.1 Global Left Ventricular Function

Figure 4A represents the variation of normalized LV volume vs time for all 20 subjects analyzed. The data show that subjects who were classified as normal, experienced similar ventricular contraction and relaxation, and their function was distinctly greater than those classified as abnormal. Accordingly, the cohort of normal subjects had a significantly higher EF (72% ± 4%) than the cohort of abnormal subjects (26% ± 12%, p < 0.00001; Fig. 4B).

**Fig. 4.**
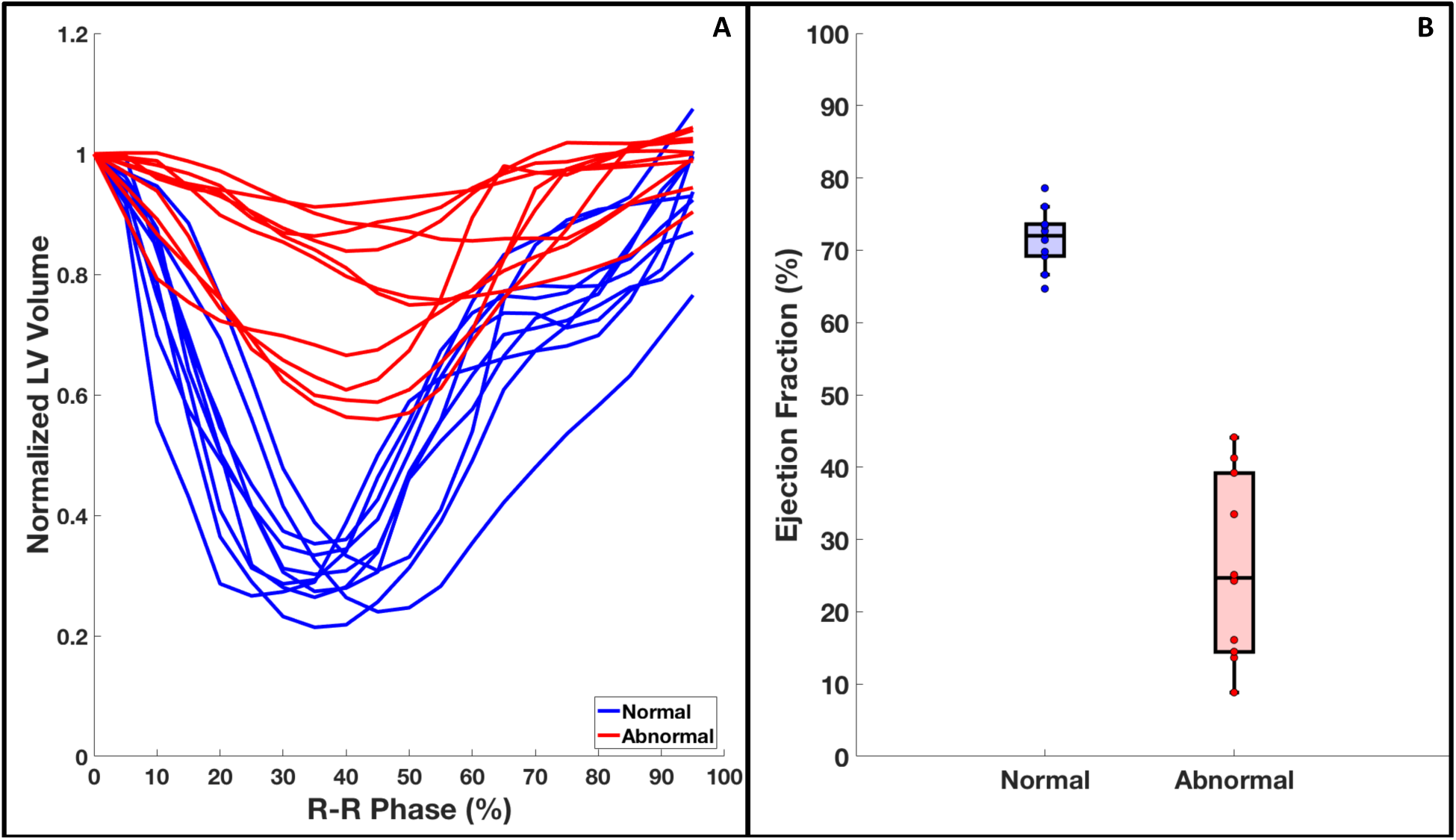
Global LV function of all 20 study subjects. (**A**) Normalized LV volume vs time across the R-R cycle for all subjects. (**B**) Boxplot showing the range of EF for the two cohorts. The normal cohort had a significantly higher EF than the abnormal cohort (72% ± 4% vs 26% ± 12%, p < 0.00001). EF: ejection fraction.

### 3.2 Regional Left Ventricular Function

Figure 5 shows the variation in endocardial topography across 6 phases of the cardiac cycle in representative short axis slices of the normal subject shown in column A of Fig. 2. The pixel colors represent the regional values of F_D_ of the endocardium. The data show a uniform decrease in topographical complexity of the base (Fig. 5A), mid-cavity (Fig. 5B), and apex (Fig. 5C) regions of the LV during systolic contraction, followed by an equivalent increase during diastolic relaxation. The regional values of F_D_ reflect these variations in the surface topography of the endocardium, supporting the hypothesis that the endocardial surface becomes smoother during systole due to the combined effect of the convergence of the trabeculae carneae and the finite resolution of the scanner.

**Fig. 5.**
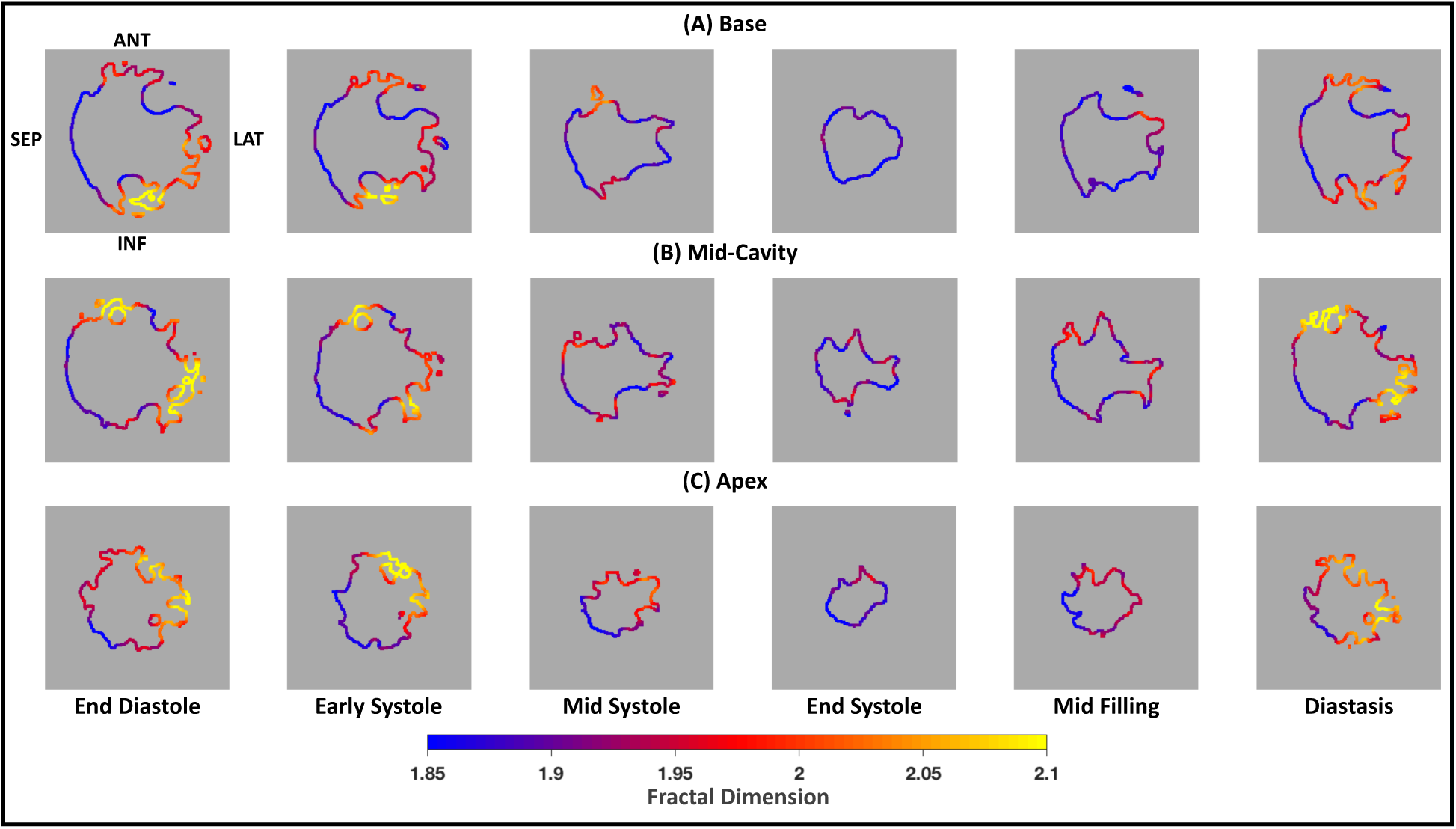
Variations in complexity of a normal LV endocardial surface across the cardiac cycle. Short axis slices of a representative normal subject (EF 73%) at three different regions (A-Basal, B-Mid, C-Apex) of the LV endocardium across 6 phases of the cardiac cycle. The regional values of F_D_ are represented by the pixel colors in the image. The orientation of the endocardium is anterior wall at 12:00, lateral wall at 3:00, inferior wall at 6:00, and septum at 9:00. All three regions showed a uniform decrease in topographical complexity of the endocardium during systole.

In contrast, Fig. 6 and 7 show similar sets of endocardial short axis slices for two representative abnormal subjects, one exhibiting regional dysfunction and the other exhibiting global dysfunction. The subject shown in Fig. 6 is the same as the subject shown in column B of Fig. 2, having an EF of 45%. The regional values of F_D_ reveal hypokinesia in the mid-cavity region (Fig. 6B), where F_D_ remains high-valued across the cardiac cycle, particularly at end-systole. However, the F_D_ values are consistent with normal function at the apex (Fig. 6C).

**Fig. 6.**
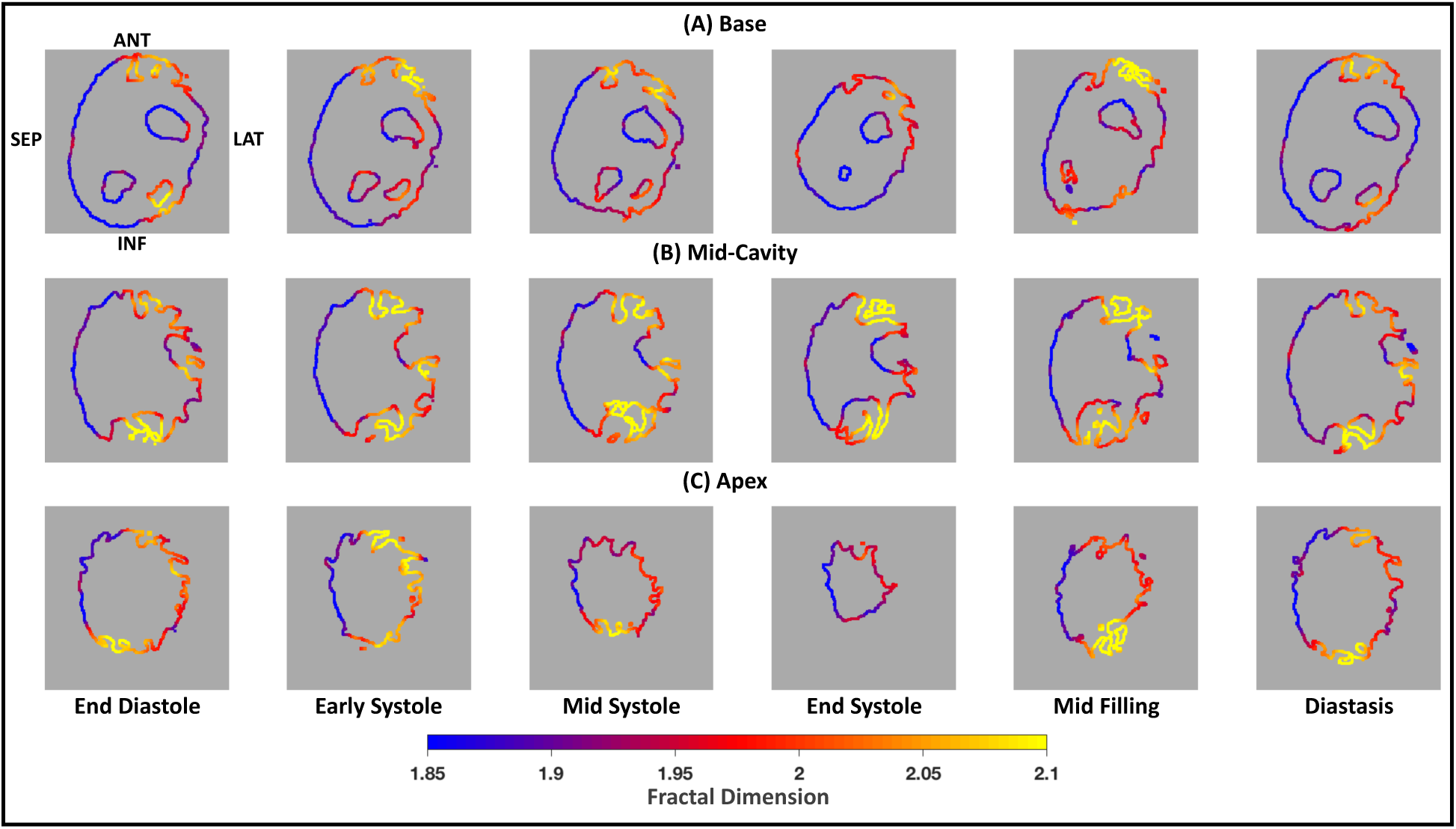
Variations in complexity of a regionally hypokinetic LV endocardial surface across the cardiac cycle. Short axis slices of a representative abnormal subject with regional LV dysfunction (EF 45% and a hypokinetic mid-cavity) at three different regions (A-Basal, B-Mid, C-Apex) of the LV endocardium across 6 phases of the cardiac cycle. The regional values of F_D_ are represented by the pixel colors in the image. The orientation of the endocardium is anterior wall at 12:00, lateral wall at 3:00, inferior wall at 6:00, and septum at 9:00. Panel B showed abnormally low variation of F_D_ across the cardiac cycle, consistent with mid-cavity hypokinesia.

**Fig. 7.**
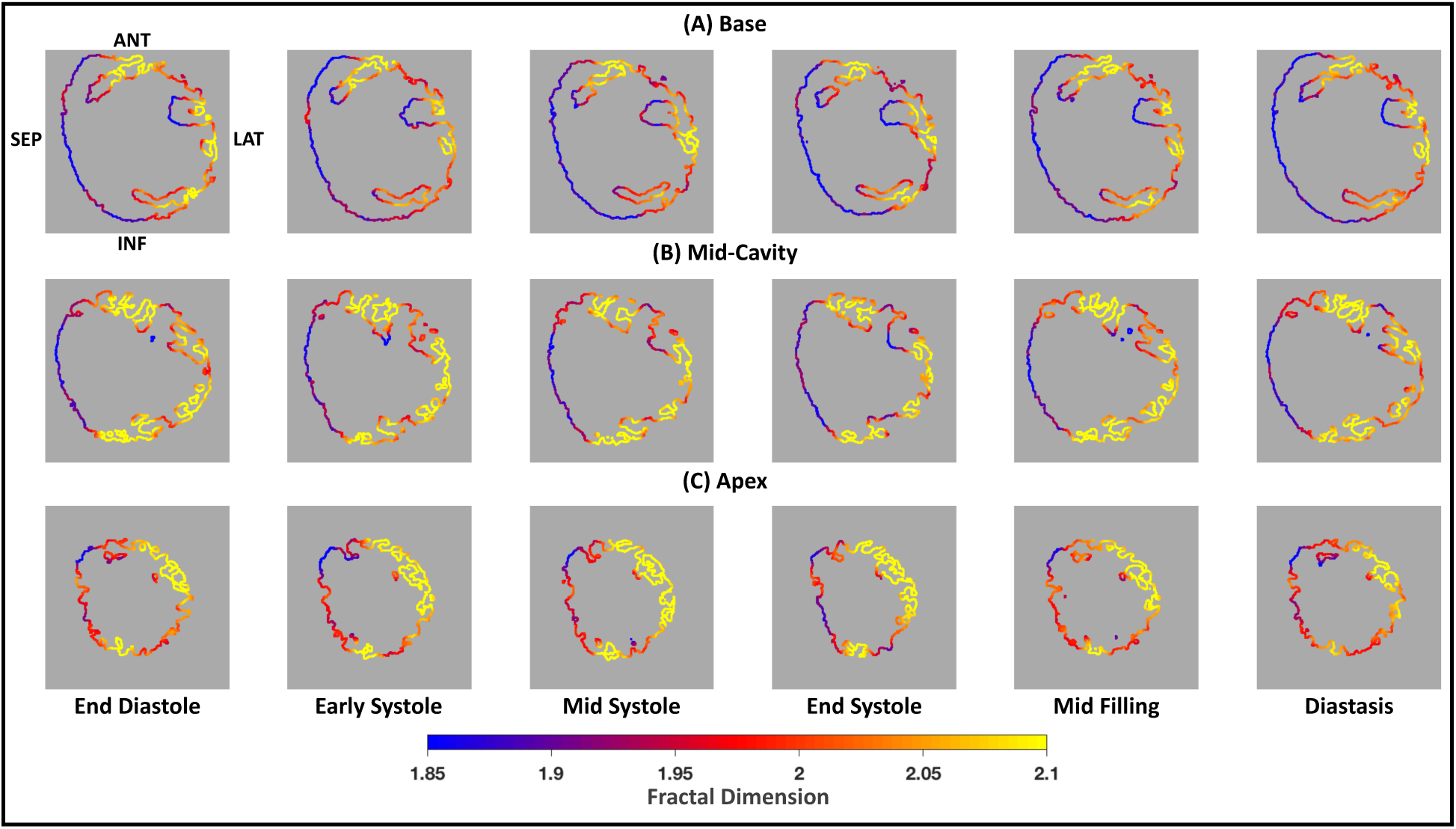
Variations in complexity of a globally hypokinetic LV endocardial surface across the cardiac cycle. Short axis slices of a representative abnormal subject with global LV dysfunction (EF 17%) at three different regions (A-Basal, B-Mid, C-Apex) of the LV endocardium across 6 phases of the cardiac cycle. The regional values of F_D_ are represented by the pixel colors in the image. The orientation of the endocardium is anterior wall at 12:00, lateral wall at 3:00, inferior wall at 6:00, and septum at 9:00. All three regions showed no significant change in topography, consistent with the fact that the subject had globally suppressed function.

The subject shown in Fig. 7 has global LV dysfunction and is shown in column C of Fig. 2, having an EF of 17%. Consistent with this subject’s global LV dysfunction, the F_D_ analysis found a lack of topography variation across the cardiac cycle in all LV regions. In fact, the maps of endocardial F_D_ at the LV base (Fig. 7A), mid-cavity (Fig. 7B), and apex (Fig. 7C) were almost identical between consecutive time frames.

Figure 8 displays temporal profiles of the spatially-averaged F_D_ in all 16 AHA segments for the 20 subjects analyzed. The F_D_ profiles of the abnormal subjects showed smaller changes during the cardiac cycle than the normal subjects in all free-wall segments. Specifically, they did not follow the normal trend consisting of a marked systolic decrease in F_D_ followed by an increase during diastolic relaxation; instead, F_D_ remained elevated with modest temporal fluctuations in the abnormal cases. The F_D_ in the septal segments, however, did not reflect this trend between the normal and abnormal subjects. This was due to the lack of trabecular tissue on the septal wall; hence texture variation across the cardiac cycle was minimal. These data suggest that it may be possible to use the time profiles of regional endocardial F_D_ as surrogate markers of free-wall LV function.

**Fig. 8.**
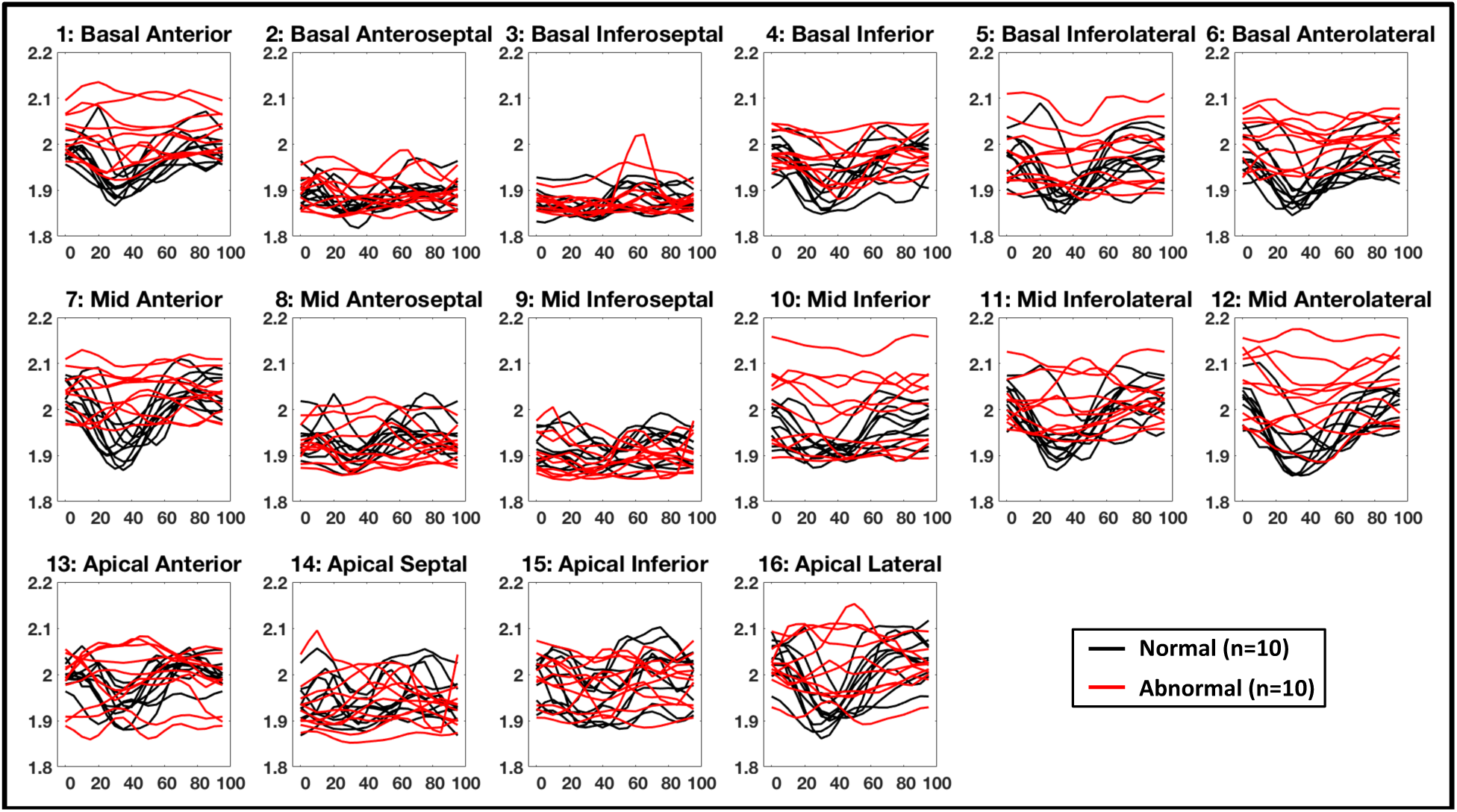
Regional F_D_ vs %R-R in all 16 AHA segments for all 20 subjects. The normal subjects showed a characteristic decrease in F_D_ in the free-wall segments during systole, while the F_D_ in the abnormal subjects remained elevated with modest temporal fluctuations. AHA: American Heart Association.

Additionally, the method can be performed using only the end-diastolic and end-systolic time frames to obtain functional estimates, thereby reducing patient exposure to radiation. To illustrate this point, Fig. 9 shows the mean percentage change in ΔF_D_ values for normal and abnormal subjects in all free wall segments computed from just the end-diastolic and end-systolic time frames. The values for the normal and abnormal cohorts were significantly different (5.9% ± 2% vs 2% ± 1.2%, p < 0.00001). Full R-R image data was used in this study to demonstrate the reproducible nature of the measurements.

**Fig. 9.**
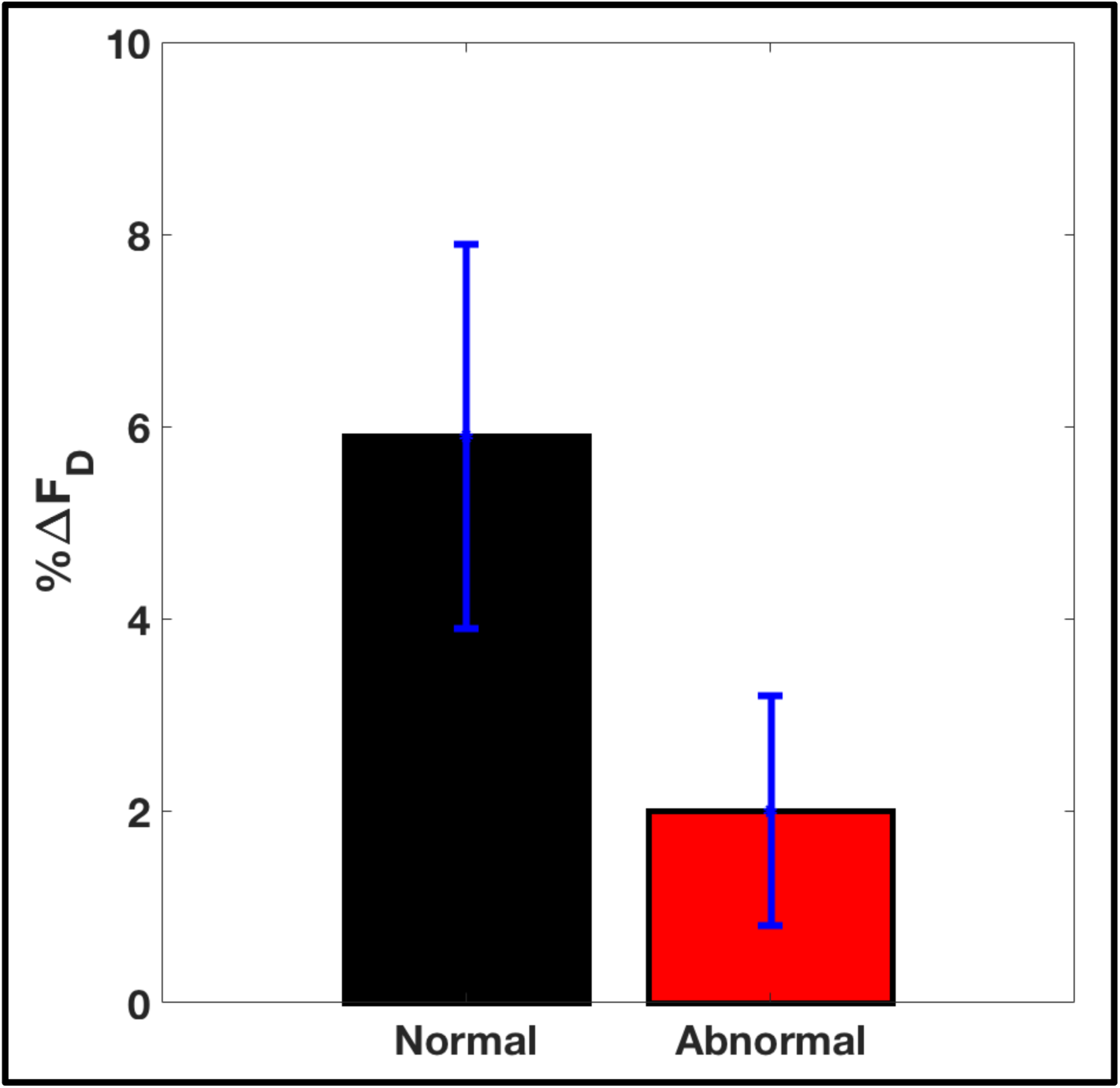
Mean peak systolic change in F_D_ (ΔF_D_) between the end-diastolic and the end-systolic phases in all free wall segments for all 20 subjects. Normal and abnormal subjects showed statistically different percentage changes in ΔF_D_ (5.9% ± 2% vs 2% ± 1.2%, p < 0.00001). AHA: American Heart Association.

### 3.3 Comparison between ε_rr_ and ΔF_D_

Peak end-systolic radial strain (ε_rr_) was measured in all 16 AHA segments for all subjects used in the study. Figure 10 plots ε_rr_ against the corresponding value of ΔF_D_ for each subject in all 16 AHA segments. Even though the two metrics measure fundamentally different physical quantities, in the small number of subjects, ε_rr_ and ΔF_D_ showed a consistent relationship in the free-wall segments. A ΔF_D_ threhold of 0.07 (vertical dashed blue lines in Fig. 10) differentiated normal from abnormal function with an accuracy of 86% in the 11 non-septal segments.

**Fig. 10.**
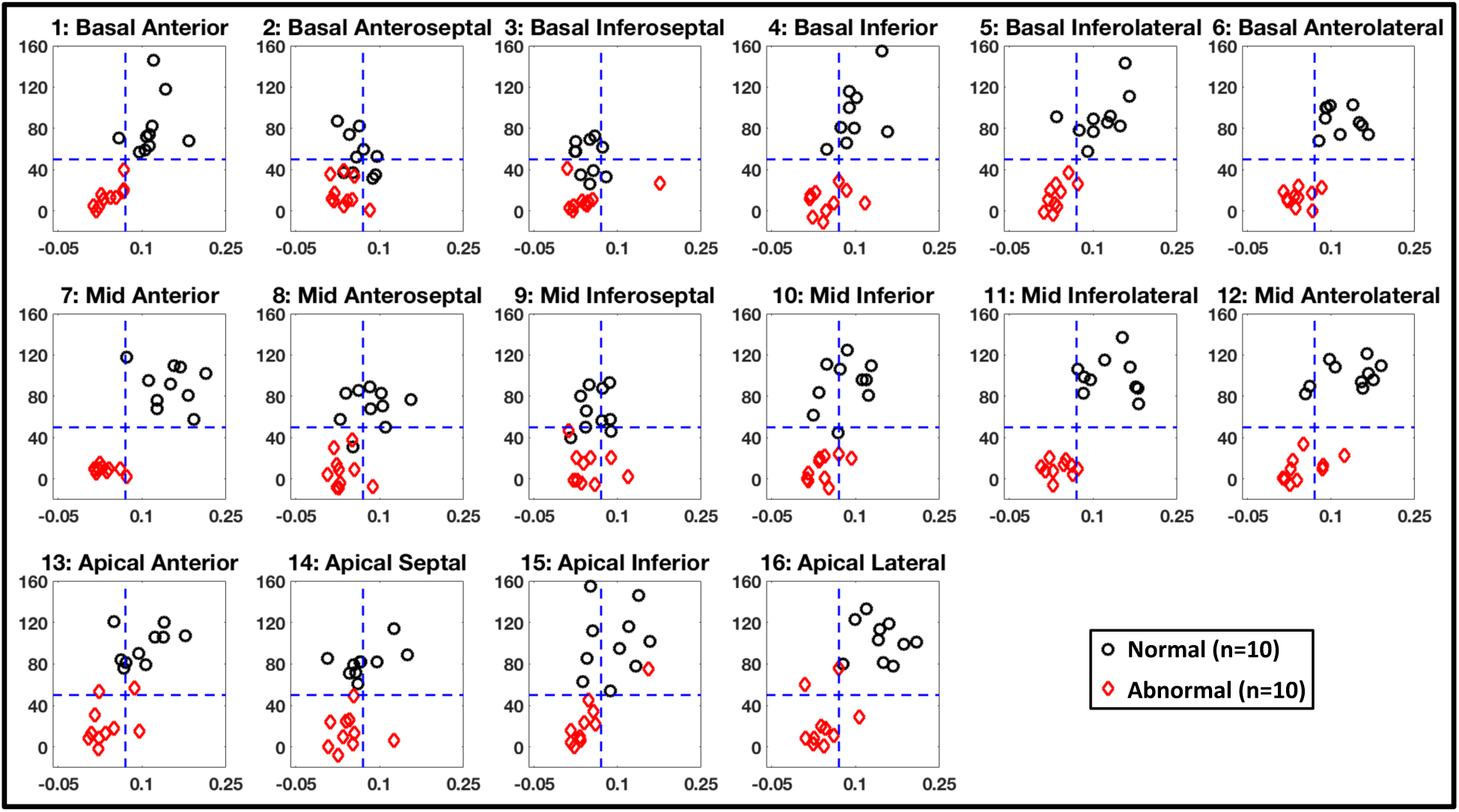
Correlation between peak systolic radial strain (εrr, shown as % on the y-axis) and peak systolic change in F_D_ (ΔF_D_, shown on the x-axis) in all 16 AHA segments for all 20 subjects used in the study. The dashed blue lines show the thresholds for ε_rr_ (50%; horizontal) and ΔF_D_ (0.07; vertical) to differentiate normal from abnormal function. The ε_rr_ threshold value of 50% was obtained from literature^20^; the ΔF_D_ value was the threshold calculated that optimized the accuracy (86%) of the test in the 11 non-septal segments. AHA: American Heart Association.

### 3.4 Comparison between Temporal Profiles of RS and Temporal Profiles of F_D_

The data presented above suggests that unsupervised analysis of the main features of the regional *F*_*D*_*(t)* profiles might provide a means for differentiating normal from abnormal wall motion. To provide proof of concept of this idea, a principal component analysis (PCA) was performed on the *F*_*D*_*(t)* profiles. Figure 11 (top panel) shows the PCA analysis on the *F*_*D*_*(t)* profiles in a sample free-wall AHA segment. As described in Sec. 2.7, only the first two PCA modes were retained for simplicity (Fig. 11, column B) and when the weights of the first mode (a_1_) were plotted against those of the second mode (a_2_) for all subjects, the normal and abnormal subjects clustered on opposite sides of the *a*_*1*_ *+ a*_*2*_ *= 0* diagonal (Fig. 11, column C). This result implies that uncomplicated data compression algorithms might be developed for unsupervised classification of normal and abnormal function from *F*_*D*_*(t)*.

**Fig. 11.**
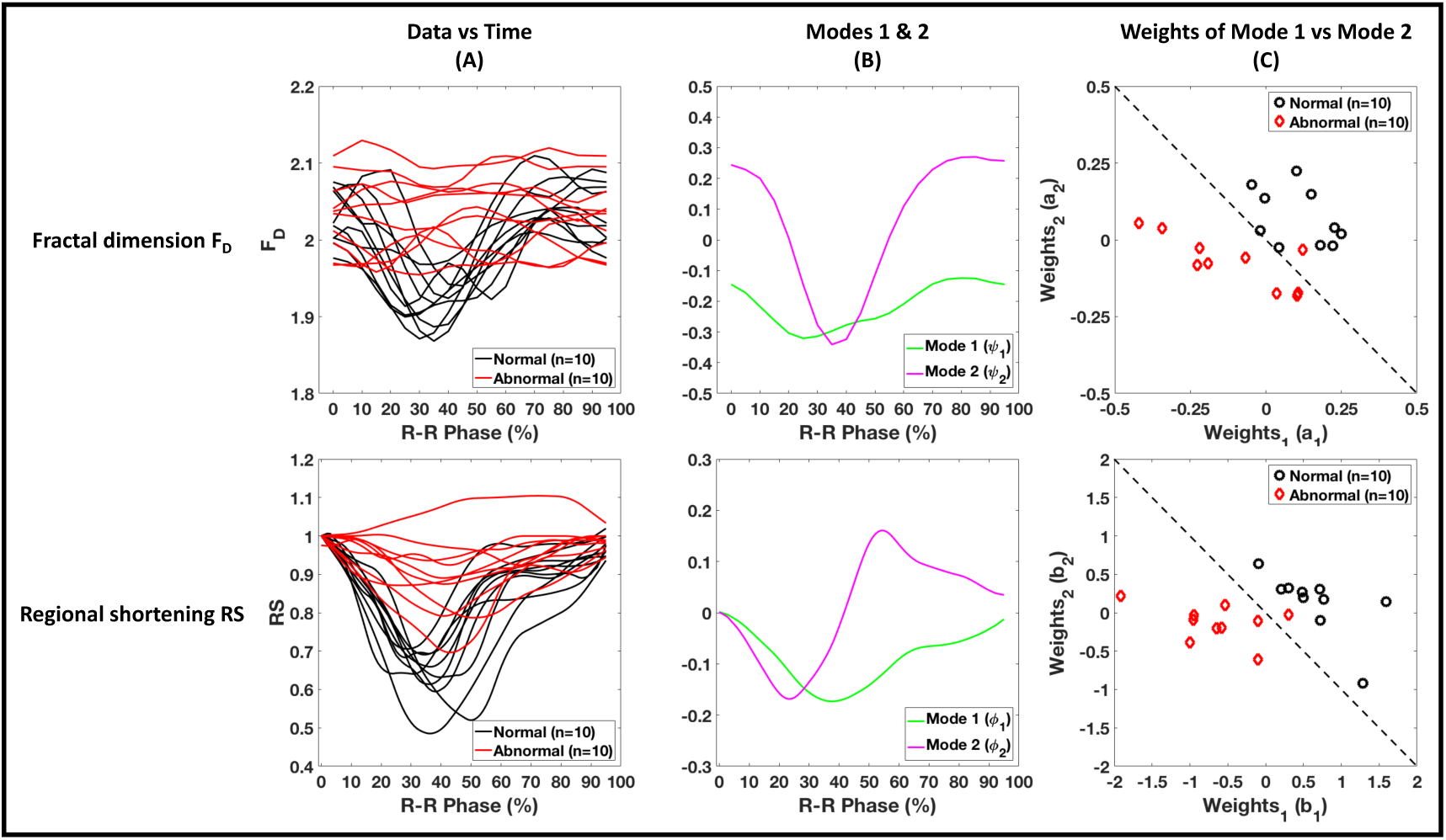
Principal component analysis on the regional values of F_D_(t) (top panel) and RS(t) (bottom panel) in a sample free-wall AHA segment for all 20 subjects. (**A**) Data (F_D_ and RS) vs time across the cardiac cycle for all subjects. (**B**) The first two modes (components) of the analysis as a function of time. (**C**) Weights of mode 1 plotted against the weights of mode 2 for all subjects. AHA: American Heart Association.

Additionally, to put this result into context, a PCA was performed in an identical manner on the *RS(t)* profiles. Figure 11 (bottom panel) shows the PCA analysis on the *RS(t)* profiles in the same sample AHA segment as for the *F*_*D*_*(t)* profiles. Again, when the weights of mode 1 (b_1_) were plotted against those of mode 2 (b_2_) for all subjects, the normal and abnormal subjects clustered on opposite sides of the *b*_*1*_ *+ b*_*2*_ *= 0* diagonal, in a similar manner to the PCA components of the regional *F*_*D*_*(t)* profiles. Table 1 summarizes the PCA results performed on the *F*_*D*_*(t)* and *RS(t)* profiles for all 16 AHA segments. All metrics were calculated based on the secondary diagonal differentiating the weights of mode 1 plotted against those of mode 2; subjects falling in the bottom left triangle were classified as true positive and subjects falling in the top right triangle were classified as true negative.

**Table 1.** Summary of the PCA on the *FD(t)* and *RS(t)* profiles for all 16 AHA segments in all 20 subjects. All metrics were calculated based on the secondary diagonal differentiating the weights of mode 1 when plotted against those of mode 2 of the PCA. Subjects on the bottom left of the secondary diagonal were classified as true positive and those on the top right of the secondary diagonal were classified as true negative. Septal segments, lacking trabeculation are italicized. AHA: American Heart Association, PPV: Positive predictive value, NPV: Negative predictive value.

| Segment # | F<sub>D</sub> |  |  | RS |  |  |
| --- | --- | --- | --- | --- | --- | --- |
|  | *Accuracy* | *PPV* | *NPV* | *Accuracy* | *PPV* | *NPV* |
| 1 | 85 | 89 | 82 | 100 | 100 | 100 |
| 2 | 65 | 71 | 62 | 95 | 91 | 100 |
| 3 | 45 | 44 | 46 | 95 | 91 | 100 |
| 4 | 65 | 67 | 64 | 95 | 91 | 100 |
| 5 | 75 | 86 | 70 | 100 | 100 | 100 |
| 6 | 90 | 100 | 83 | 100 | 100 | 100 |
| 7 | 95 | 100 | 91 | 95 | 100 | 91 |
| 8 | 55 | 57 | 54 | 90 | 90 | 90 |
| 9 | 40 | 40 | 40 | 95 | 91 | 100 |
| 10 | 80 | 100 | 71 | 95 | 91 | 100 |
| 11 | 75 | 86 | 69 | 100 | 100 | 100 |
| 12 | 95 | 100 | 91 | 100 | 100 | 100 |
| 13 | 70 | 75 | 67 | 95 | 100 | 91 |
| 14 | 60 | 63 | 58 | 95 | 91 | 100 |
| 15 | 55 | 55 | 56 | 95 | 100 | 91 |
| 16 | 75 | 86 | 69 | 100 | 100 | 100 |

### 3.4 Reproducibility and Threshold Sensitivity

To demonstrate the reproducibility of the regional F_D_ estimates, Fig. 12 shows each independently calculated F_D_ value at a specific instant of the cardiac cycle overlaid onto the temporally smoothed *F*_*D*_*(t)* profiles in a sample free-wall AHA segment for all 20 subjects in the study. Each time frame was analyzed independently and every third image in these sequences had no views shared in the image reconstruction; therefore, the noise was independent. The F_D_ vs time profiles across the cardiac cycle show the natural variability of the data under the assumption that the fractal dimension will evolve with a smooth pattern in time. Figure 12 also highlights the fact that ΔF_D_ is much larger than the variance in independent F_D_ estimates around the smooth curve.

**Fig. 12.**
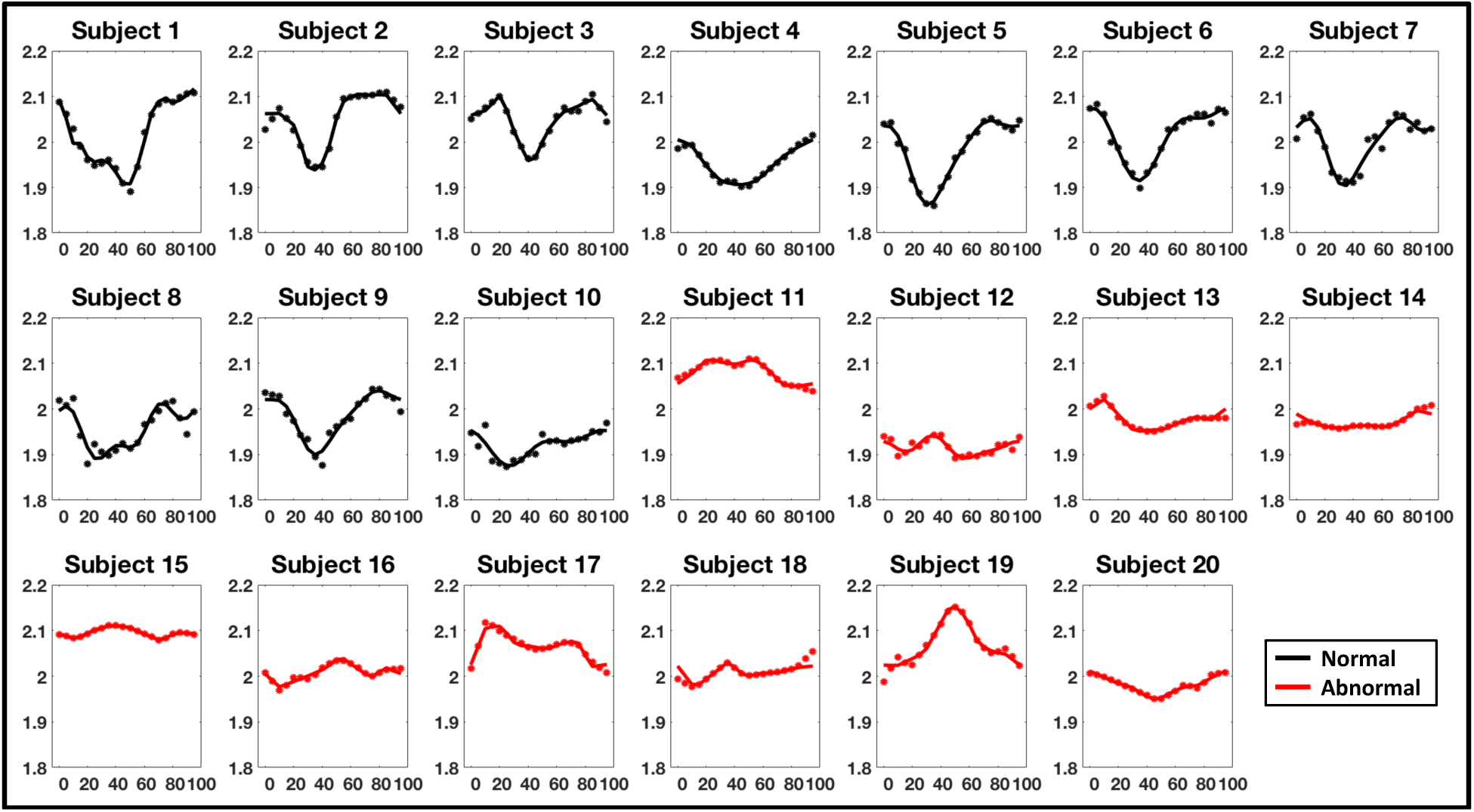
Reproducibility in measurements of regional F_D_ estimates vs %R-R in a sample free-wall AHA segment for all 20 subjects. Independent F_D_ estimates (dots) calculated at each instant of time of the cardiac cycle overlaid onto the temporally smoothed F_D_(t) profiles. Variance in the independent F_D_ estimates around the temporally smoothed curves provides an estimate of the reproducibility. AHA: American Heart Association.

Figure 13 shows the sensitivity of ΔF_D_ to the choice of the user input threshold for LV blood volume segmentation. Over the domain of reasonable threshold values for the LV blood volume, the absolute F_D_ estimate will depend on the choice of threshold; hence the change in F_D_ between end-diastole and end-systole (ΔF_D_) was used to investigate the sensitivity. Also, currently ΔF_D_ is used as the biomarker for differentiating normal from abnormal wall motion. For two sample normal and two sample abnormal subjects, the LV was segmented for thresholds in the range of 100 to 1000 HU. Each subject had a lower limiting threshold value, below which other structures such as the myocardium and the RV were included. Similarly, each subject had an upper limiting threshold value beyond which the LV disintegrated rapidly. Within these limits of relatively stable LV volumes, ΔF_D_ was calculated in a sample free-wall AHA segment for each subject. Figure 13A plots the end-systolic volume of the segmented blood pool as a function of threshold and Fig. 13B plots the variation in ΔF_D_ within the stable threshold regime (marked with black dots in Fig. 13A). The ΔF_D_ values remained very stable over a wide range of 175 HU, which is well beyond the range of the variation in threshold obtained by the user from the process described in Eq. 1. Figures 13C and 13D plot F_D_(t) profiles across the cardiac cycle for normal Subject 1 (Fig. 13C) and for normal Subject 2 (Fig. 13D) for three different threshold values sampling the range of stable threshold values for blood segmentation. While the absolute value of F_D_ changes as a function of threshold, the temporal curves of both subjects for all three threshold values show a characteristic decrease in F_D_ during systolic contraction followed by an increase during diastolic relaxation, highlighting the stability in detecting regional wall motion across the cardiac cycle.

**Fig. 13.**
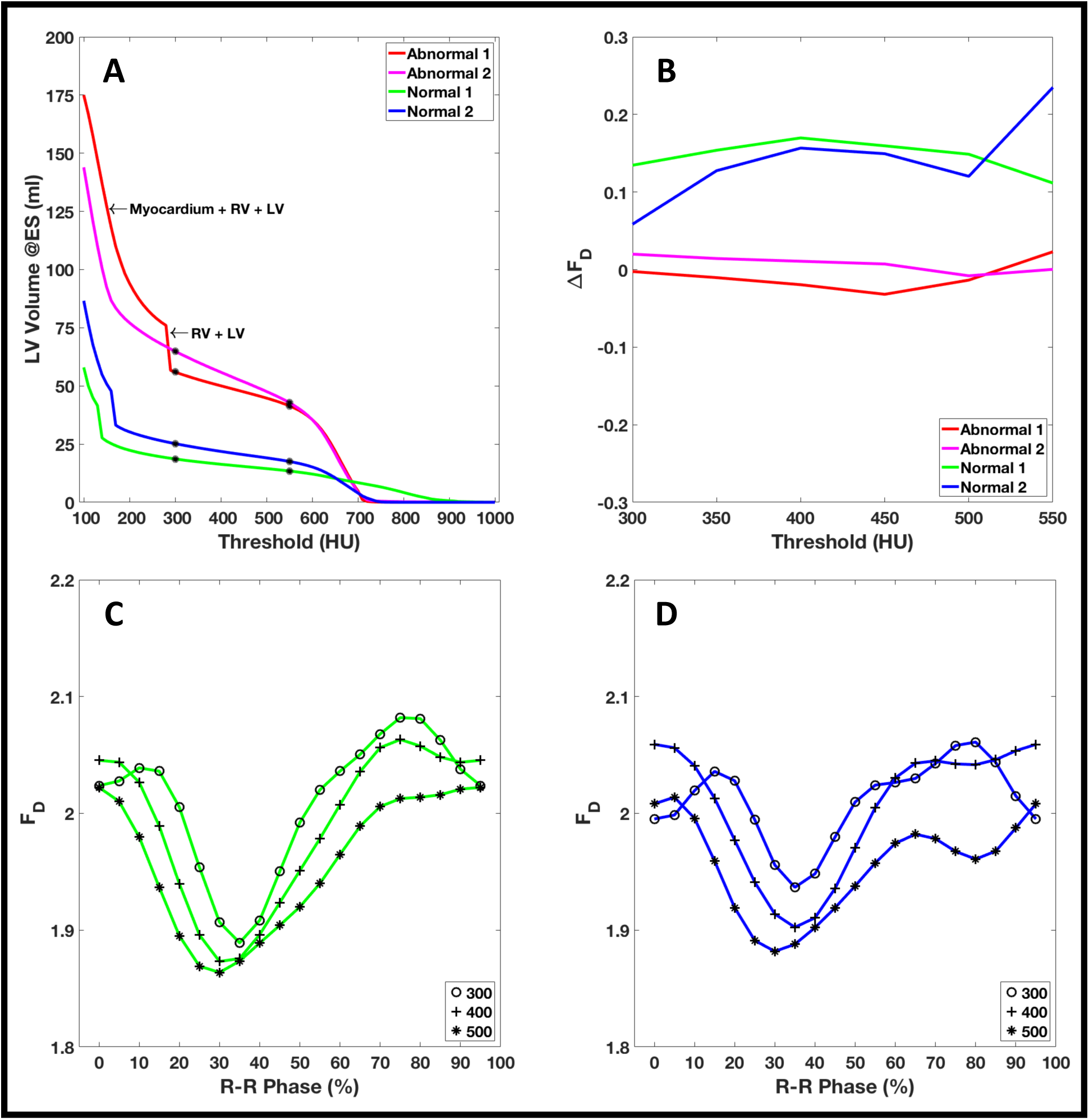
Threshold sensitivity analysis. (**A**) LV volume at end-systole as a function of threshold for two sample normal and two sample abnormal subjects. (**B**) Variation of ΔF_D_ in the regime of stable threshold values for each subject in a sample free-wall AHA segment. The stable threshold regime is marked with black dots in (A). (**C**) F_D_(t) profiles across the cardiac cycle for normal Subject 1 for three different threshold values sampling the stable threshold regime. (**D**) F_D_(t) profiles across the cardiac cycle for normal Subject 2 for three different threshold values sampling the stable threshold regime. AHA: American Heart Association.

## 4. Discussion

This manuscript presents an image analysis method that exploits the temporal changes of LV endocardial topography across the cardiac cycle to quantify regional left ventricular function. The method uses scans that can be routinely acquired as part of coronary CT angiography (CCTA) studies or cardiac function assessment, exposing the patient to no additional radiation. The analysis is fast, with fractal computation times of ∼1 minute for 20 time frames running on a personal computer. Of note, the calculation of F_D_ does not involve image registration. This feature is of particular practical interest because the non-rigid deformation of the heart and the lack of a significant number of anatomical landmarks, complicate cardiac image registration^24^. Additionally, fractal dimension analysis does not require manual contours to be drawn to define the LV endocardium and epicardium, which can be susceptible to inter and intra-observer variability^25^. With current advances in automatic cardiac segmentation^26–29^, it is reasonable to expect fractal dimension analysis to become fully automated. Since the method is also independent of motion tracking, it can be used to characterize topography even on a single static image of the heart. The proposed analysis may be additionally applicable to the quantification of LV non-compaction^19^ (LVNC). Since LVNC is much less prevalent when compared to LV dysfunction from heart failure or ischemia, we did not study this group of patients. Another possible application of fractal dimension is the quantification of left atrial appendage (LAA) function, which currently lacks a robust quantitative method. Additionally, we are also encouraged by the fact that novel motion-estimation and motion-compensated reconstruction algorithms^30^ will only improve the fidelity of trabeculae seen on the CT images. Previous studies have measured the average fractal dimension over the LV; the work reported here relates changes in regional F_D_ values of the LV endocardium to local myocardial function for the first time. This work also demonstrates that previously reported average values of fractal dimension ignore the fact that the smooth surface of the septum is quite distinct from the free wall.

The image sequences used in this study were obtained using CT scanner manufacturers’ standard reconstruction algorithms as implemented clinically. Images spanning the complete cardiac cycle were acquired over a single heartbeat, which is a significant advantage over other imaging modalities requiring multiple beat acquisitions to produce a 3D volume. Echocardiography is the most widespread modality to assess regional cardiac function in the clinical setting. However, it depends on the skill of the sonographer^31^ and often involves the semi-quantitative visual assessment of regional wall motion^32,33^, leading to significant inter and intra-observer variability^34,35^. Moreover, echocardiographic 2D assessment of wall motion may not fully capture the complicated 3D motion of the heart, and may be affected by tissue entering and leaving the imaging plane^36^. CMR tissue tagging is the current gold standard in the assessment of regional cardiac function^37–40^. However, its use has primarily remained in the research setting due to the requirements of extended breath holding and multiple heartbeat acquisitions, together with its limited resolution^41^. Additionally, the application of CMR in patients implanted with some metallic devices, particularly cardiac assisting devices (pacemakers, LVADs, etc.), is limited.

Although only a small cohort of normal and abnormal subjects was used in this study, as a proof of concept, our results highlight the ability of fractal dimension analysis to differentiate between normal and abnormal free-wall motion by evaluating the changes in regional values of F_D_ across the cardiac cycle. Reproducible F_D_ estimates were obtained which were stable across a wide range of user input thresholds for LV blood segmentation. The method can obtain functional estimates from just the end-diastolic and end-systolic phases of the cardiac cycle, thereby reducing patient exposure to ionizing radiation. Furthermore, we found that ΔF_D_ correlated well with peak end-systolic radial strain in the free-wall segments of the LV. Finally, we demonstrated as a proof of principle that the fractal dimension vs. time data can be analyzed using simple well known tools such as principal component analysis to potentially classify subjects as normal or abnormal based on regional LV function.

### 4.1 Limitations

While it is clear that wall motion across the cardiac cycle leads to temporal changes in endocardial F_D_, an explicit relationship between wall strain and F_D_ is yet to be established. The trabeculae in the heart are irregular in structure and vary between individuals^42,43^, which makes it challenging to derive baseline values of normal and abnormal function. The septal wall has little trabeculation, especially near the LV outflow tract; therefore, its topography remains smooth with small changes in F_D_ across the cardiac cycle, even in ventricles with normal function. Thus, fractal dimension analysis will likely only be applicable to non-septal regions.

While we have shown in this study that fractal dimension analysis quantifies regional LV function that correlated well with radial strain and with endocardial regional shortening measurements from CT, we did not perform a head-to-head validation in patients with CMR tagging; the gold-standard imaging modality for strain. The promising initial results presented here motivate future studies to address validation in larger numbers of patients and to determine the extent of the LV for which fractal dimension analysis can be used.

Currently, our image processing pipeline requires the user to input a threshold value for LV blood volume segmentation. While this is not ideal, we note that the aim of this work was to evaluate the feasibility of using topography variation as a surrogate measure of cardiac function, rather than to develop an optimized segmentation algorithm. Given the current advances in machine assisted segmentation techniques, such as convolutional neural networks^26–29^, we expect that the choice of threshold will very shortly not be subject to inter-user variability.

Low x-ray dosage implies low SNR of the images, which in turn has the potential to affect the imaged topography of the endocardium, especially if it introduces the need for additional smoothing during CT image reconstruction. Thus, dose modulated scans might offer inconsistent topography across the cardiac cycle, which could introduce artifacts in the temporal changes of F_D_. Future work should address the effects of dose and dose modulation on the computed values of fractal dimension.

While we have demonstrated the applicability of the method to images acquired with standard clinical protocols, the variability in CT image acquisition (different scanner models) and CT image reconstruction (different fields of view, slice thicknesses, and reconstruction kernels) of these images may introduce biases in the fractal dimension estimates. A study investigating the independent effects of these scan parameters needs to be undertaken.

## Appendix A: Validation of the box-counting algorithm

The box-counting algorithm developed in-house, which was implemented in the FORTRAN programming language, was validated on synthetically created images of standard fractals well known in the literature. Three Euclidean 2D synthetic fractals (Julia Set (c=-1), Julia Set (c=-0.123+0.745i), and Sierpinski Triangle) and five Euclidean 3D synthetic fractals (3D Cantor Cube, 3D Random Cantor Dust (p=0.5), 3D Random Cantor Dust (p=0.6), 3D Random Cantor Dust (p=0.7), and Menger Sponge) were created, as shown in Fig. 14. Table 2 summarizes the resolutions, the theoretical and computed fractal dimensions, and the percent error between the two values for each test fractal.

**Table 2.**
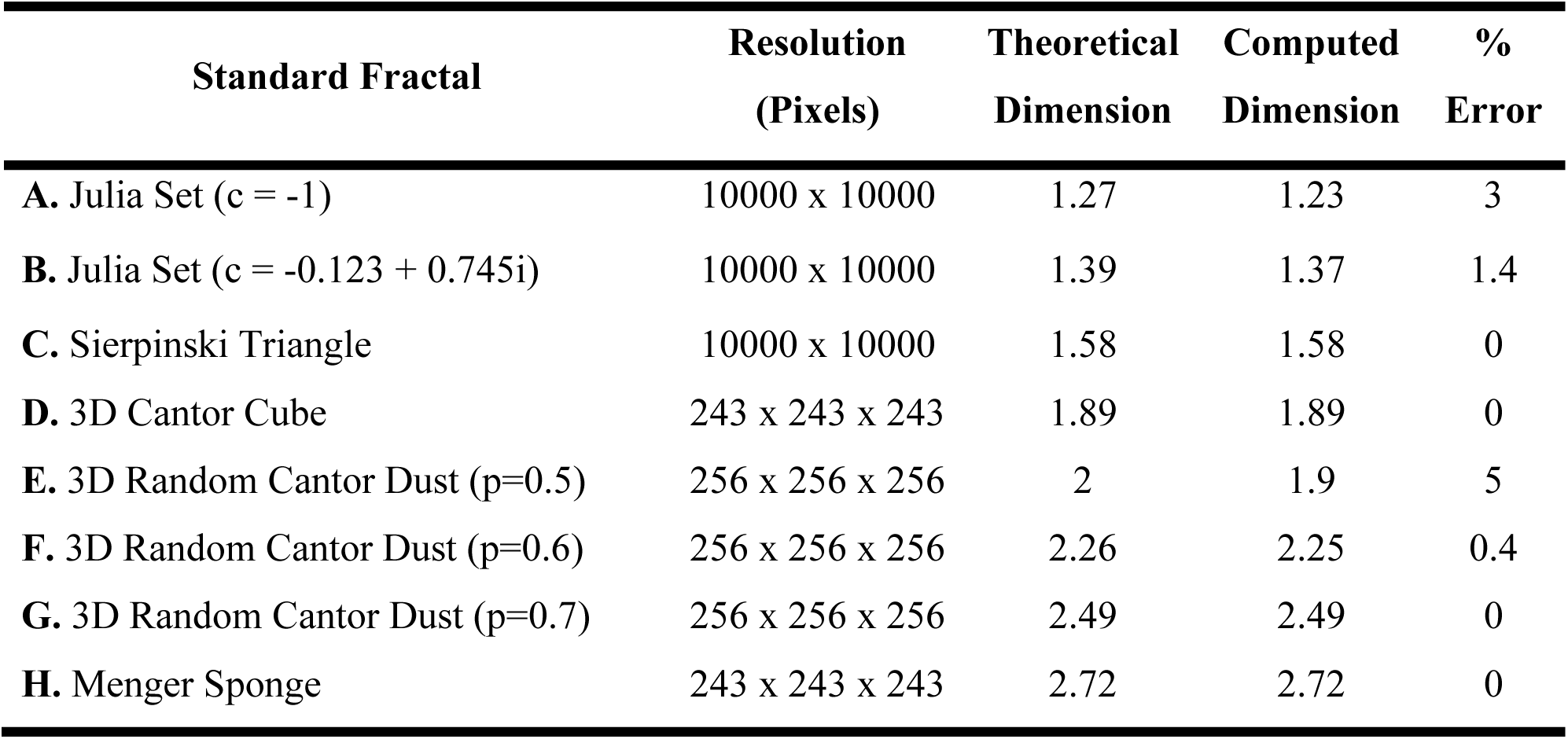
Summary of the theoretical and computed fractal dimensions for the 8 synthetic standard fractal images.

**Fig. 14.**
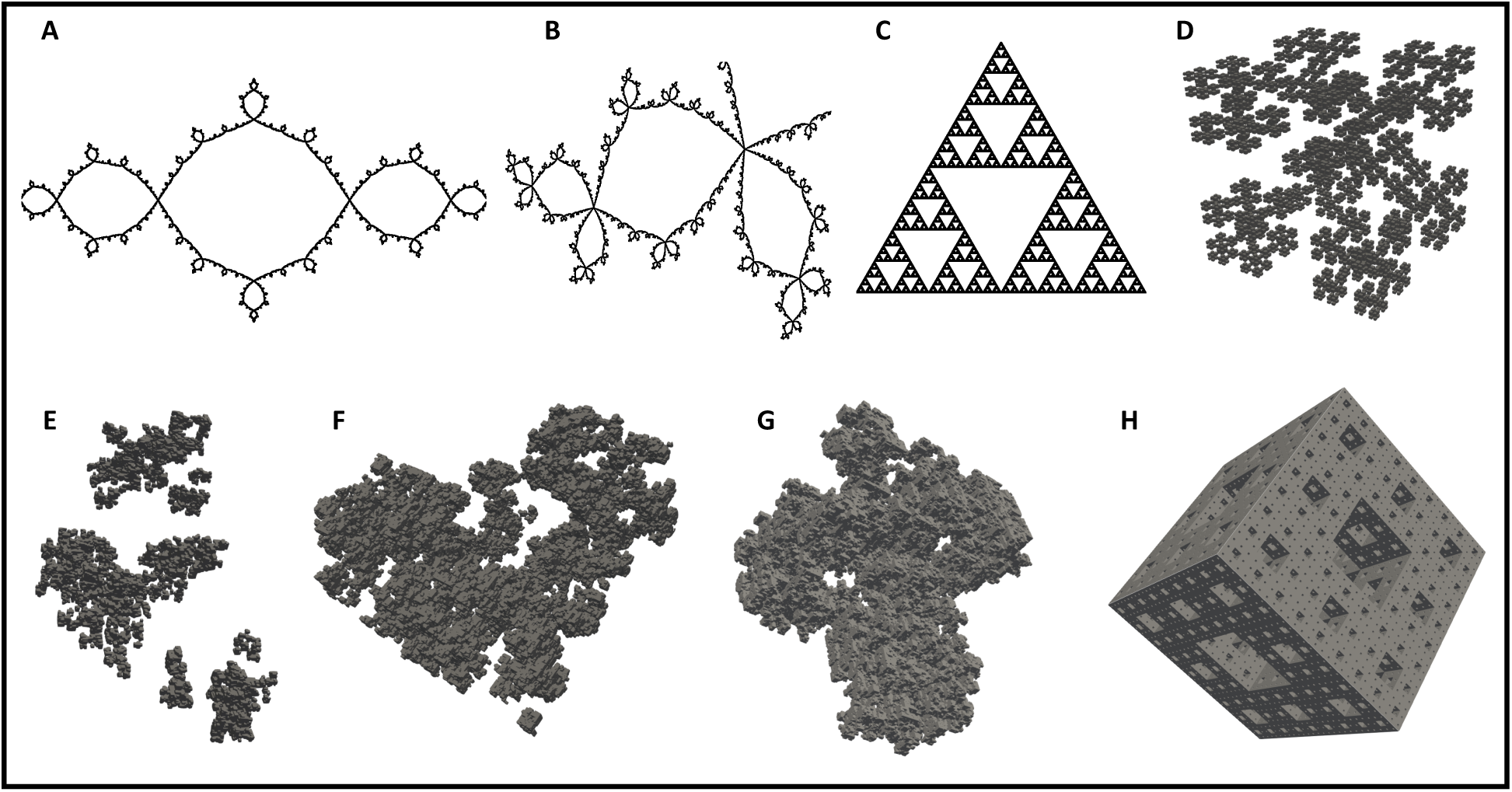
Theoretical fractals used for validating the in-house developed box-counting algorithm. (**A**) Julia Set (c=-1). (**B**) Julia Set (c=-0.123+0.745i). (**C**) Sierpinski Triangle. (**D**) 3D Cantor Cube. (**E**) 3D Random Cantor Dust (p=0.5). (**F**) 3D Random Cantor Dust (p=0.6). (**G**) 3D Random Cantor Dust (p=0.7). (**H**) Menger Sponge.

## Acknowledgements

This work was funded in part by the University of California President’s Postdoctoral Fellowship Program (Francisco Contijoch) and the NIH grants T32 HL105373 “Integrative Bioengineering of Heart, Vessels, and Blood” (Gabrielle Colvert) and R01 HL144678 (Elliot McVeigh).

**Ashish Manohar** received a BE from the R.V. College of Engineering, India and an MS from UC San Diego, USA, both in mechanical engineering. He joined the Cardiovascular Imaging Laboratory (CViL) in 2017, where he is currently pursuing his doctoral degree in mechanical engineering. His research interests include cardiac function assessment from CT images, and understanding the spatial and temporal resolution limits of 4DCT in the estimation of regional wall motion abnormalities.

**Elliot McVeigh**, BSc in Physics and PhD in Medical Biophysics from University of Toronto. Faculty in BME and Radiology at Johns Hopkins from 1988-1999. PI at NIH/NHLBI from 2000-2007. From 2007 through 2015 he was Chairman of BME at JHU. Since 2015 he has been a Professor of Bioengineering, Radiology and Cardiology at UCSD. His laboratory focuses on creating novel MRI and CT techniques for diagnosis and guidance of therapy of heart disease.

